# Hybrid optogenetic and electrical stimulation for greater spatial resolution and temporal fidelity of cochlear activation

**DOI:** 10.1101/2020.07.27.187294

**Authors:** Alex C. Thompson, Andrew K. Wise, William L. Hart, Karina Needham, James B. Fallon, Niliksha Gunewardene, Paul R. Stoddart, Rachael T. Richardson

## Abstract

Compared to electrical stimulation, optogenetic stimulation has the potential to improve the spatial precision of neural activation in neuroprostheses, but it requires intense light and has relatively poor temporal kinetics. We tested the effect of hybrid stimulation, which is the combination of subthreshold optical and electrical stimuli, on spectral and temporal fidelity in the cochlea by recording multiunit activity in the inferior colliculus of channelrhodopsin (H134R variant) transgenic mice. Pulsed light or biphasic electrical pulses were delivered to cochlear spiral ganglion neurons of acutely deafened mice, either as individual stimuli or as hybrid stimuli for which the timing of the electrical pulse had a varied delay relative to the start of the optical pulse. Facilitation occurred when subthreshold electrical stimuli were applied at the end of, or up to 3.75 ms after subthreshold optical pulses. The spread of activation resulting from hybrid stimulation was significantly narrower than electrical-only and optical-only stimulation (p<0.01), measured at equivalent suprathreshold levels of loudness that are relevant to cochlear implant users. Furthermore, temporal fidelity, measured as maximum following rates to 300 ms pulse trains bursts up to 240 Hz, was 2.4-fold greater than optical-only stimulation (p<0.05). By significantly improving spectral resolution of electrical- and optical-only stimulation and the temporal fidelity of optical-only stimulation, hybrid stimulation has the potential to increase the number of perceptually independent stimulating channels in a cochlear implant.

## Introduction

Cochlear implants restore hearing percepts to people with severe or profound hearing loss. The device directly stimulates the auditory neurons with electrical pulses, thus bypassing damaged or lost sensory hair cells in the cochlea. Taking advantage of the tonotopic arrangement of the cochlea, sound is processed into up to 22 discrete frequency bands and delivered to the most relevant site in the cochlea. Ideally, the cochlear implant would also use precise timing of electrical pulses at each stimulation site to encode pitch (Erfanian Saeedi et al., 2017) and other fine timing information. In practice, however, current spread causes significant overlap between the neural populations activated by each electrode (O’Leary et al., 2009, Kral et al., 1998, Black et al., 1981, Snyder et al., 2004) making it nearly impossible to get fine temporal structure at all stimulating sites. Consequently, cochlear implants stimulate regions of the cochlea sequentially and users report difficulty understanding speech in everyday background noise and poor perception of tonal sounds such as music (Fu et al., 1998, Friesen et al., 2001, McDermott, 2004, Wilson and Dorman, 2008). To improve the representation of complex sounds, an implant must be able to provide high spectro-temporal resolution i.e. very fine control over the site and timing of neural activation.

Bipolar, tripolar or focussed multipolar stimulation strategies have improved spectral resolution in animal studies (Berenstein et al., 2008, George et al., 2015, George et al., 2014, Bierer and Middlebrooks, 2004) and human trials (Donaldson et al., 2005, Srinivasan et al., 2012, Landsberger and Srinivasan, 2009, Bierer, 2007). However, they have not delivered fine temporal structure for simultaneous multi-channel stimulation. As such, there has not been any clear clinical benefit from the use of these stimulation strategies (Berenstein et al., 2008, Bierer and Litvak, 2016). Optical stimulation of auditory neurons presents an alternative method of increasing spectral resolution as light can be focused onto a target and is not limited by the same conductive spread as electrical current (reviewed in (Richardson et al., 2020)). Activation in the cochlea with a spectral resolution approaching that of acoustic input has been achieved with infrared or near infrared light (Izzo et al., 2007, Richter et al., 2008, Guan et al., 2015, Wang et al., 2016, Richter et al., 2011). Infrared neural activation has proven controversial, with some studies suggesting photoacoustic mechanisms of activation (Thompson et al., 2015, Kallweit et al., 2016) and others indicating direct neural activation (Tan et al., 2018). Tissue heating by the infrared light is also likely to severely limit performance for closely spaced channels at high repetition rates (Thompson et al., 2013).

Using another approach, optogenetic methods, in which auditory neurons are first genetically modified with light-sensitive ion channels (opsins), has enabled neural activation with relatively low power blue or red visible light sources and has also resulted in very high spectral resolution of activation in the cochlea (Wrobel et al., 2018, Keppeler et al., 2018, Duarte et al., 2018, Hernandez et al., 2014, Mager et al., 2018, Dieter et al., 2019). In optogenetic-based stimulation, one of the main limitations is the slow temporal kinetics of the opsins used. Typical electrical stimulation rates for the cochlear implant vary, but the ACE (Advanced Combination Encoder) strategy, for example, uses fixed rates between 250 Hz and 2.4 kHz. Modulations above 300 Hz have been demonstrated to be important for pitch perception (Croghan et al., 2017, Smith et al., 2002), but despite the potential for very high rate stimulation, there is no evidence of improved speech perception when using electrical stimulation rates over 500 Hz (Shannon et al., 2011). The maximum firing rate of optically evoked action potentials is limited by the channel closing kinetics of the opsin, which for channelrhodopsin-2 (ChR2) is only 70 Hz (Hernandez et al., 2014). One strategy to overcome slow temporal kinetics is to use ultrafast opsins such as Chronos (Klapoetke et al., 2014), where a 50% spike probability was achieved with repetition rates of 200 Hz or more (Keppeler et al., 2018), but required intense light with associated power and safety issues. Another strategy is to combine subthreshold electrical pulses with subthreshold optical pulses (hybrid stimulation). Our earlier work indicated that hybrid stimulation resulted in action potentials in cultured auditory neurons with a three-fold increase in the peak firing rate compared to optical stimulation alone (Hart et al., 2020). However, spectral resolution was not addressed in that study.

Here, we compare the spectral resolution and maximum following rate of optical, electrical and hybrid stimulation in the auditory system of transgenic mice expressing the H134R variant of ChR2, which has slightly slower kinetics but generates larger photocurrents compared to wild-type ChR2 (Berndt et al., 2011). Cochleae of acutely deafened mice were implanted with a single channel electrical/optical device enabling hybrid stimulation. Electrical-only, optical-only or hybrid stimuli were applied to the cochlea at sub- and suprathreshold levels, with the timing of the electrical pulse having a varied delay relative to the start of the optical pulse in the hybrid stimuli. Spread of activation and entrainment rates were measured via a multi-channel recording array inserted into the contralateral inferior colliculus (IC) of the auditory midbrain to record multi-unit neural activity across the isofrequency laminae (George et al., 2015, George et al., 2014, Landry et al., 2013). Our results show activation of the auditory pathway by the combination of a sub-threshold optical pulse and a sub-threshold electrical pulse (hybrid stimulation). We demonstrate an extended post-stimulus facilitation period during which a sub-threshold electrical stimulus can be applied to activate the auditory pathway *in vivo*, a finding which we also confirmed at the cellular level *in vitro* by patch clamp electrophysiology. Hybrid stimulation significantly reduced the spread of activation compared to electrical stimulation alone and also resulted in a 2.4-fold increase in the maximum following rate for pulse trains, rising from 46 Hz for optical only to 111 Hz for hybrid stimulation.

## Materials and methods

### Animals

Heterozygous transgenic mice expressing ChR2-H134R in spiral ganglion neurons were derived by breeding COP4*H134R/EYFP mice (Jax strain 012569: B6;129S-Gt(ROSA)26Sor^tm32(CAG-COP4*H134R/EYFP)Hze/J^) with Cre-parvalbumin mice (Jax strain 008069: B6;129P2-Pvalb^tm1(cre)Arbr^). C57BL/6 mice were used as controls. Males and females were used at random. The use and care of the experimental animals in this study were approved by St Vincent’s Hospital (Melbourne) Animal Ethics Committee (#14-028 and #18-003) and follows the Guidelines to Promote the Wellbeing of Animals used for Scientific Purposes (2013), the Australian Code for Care and Use of Animals for Scientific Purposes (8^th^ edition, 2013) and the Prevention of Cruelty to Animals Amendment Act (2015).

### Inferior colliculus recordings

A total of ten transgenic mice (8 males and 2 females) and two wild-type mice (both female) were used with an average age of 64 days (range 35-87 days). Cochlear exposures and IC recordings were performed under gaseous general anaesthesia; 1-2% isofluorane (Zoetis, Melbourne, Australia) mixed with oxygen delivered at a flow rate of 1.5 L/min, with additional local anaesthesia (1% Lignocaine hydrochloride; 10 mg/mL; ProVet, Melbourne, Australia) applied at each surgical site. A heating pad was used to maintain core body temperature. Respiration rate was monitored over the duration of the experiment and fluids were injected subcutaneously after 2 hours.

Surgical access to the left bulla was made by a retroauricular incision and retraction of adipose and muscle tissue. The bulla was opened manually with a blade and the opening was expanded with angled forceps. The stapedial artery was removed by coagulation with a low-temperature bipolar cautery. Hair cells were inactivated by the flushing of 8-10 drops of 100 mg/mL neomycin sulfate (Sigma-Aldrich, St Louis, MO) through the punctured round window membrane whilst aspirating from the apex via a small hand-drilled hole. The cochlear surgical site was then temporarily plugged with a saline-soaked cotton ball while the IC was exposed. For the IC exposure, mice were placed in a stereotaxic frame (David Kopf Instruments, Tujunga, CA). A craniotomy was performed in the region of the intersection of the parietal and interparietal bones. The dura mater was dissected to expose the IC contralateral to the neomycin-exposed cochlea. A hybrid optical/electrical device, described below, was then positioned 1-2 mm into the cochlea through the round window membrane via a micropositioner. With the hybrid device in place, a multiunit recording array (NeuroNexus Techologies, Ann Arbor, MI), consisting of 32 recording sites spaced at 50 μm intervals over 1550 μm, each with a surface area of 177 μm^2^, was mounted on a Microdrive positioner (David Kopf Instruments, Tujunga, CA) and positioned at the surface of the exposed IC. The array was advanced along the dorsal-ventral axis of the central nucleus of the IC to a depth of approximately 2 mm from the surface as determined by visual monitoring of responses at the tip electrode to optical or electrical stimulation in the cochlea. When the recording array was confirmed to be within the IC, a 1% agar solution was applied to the surface of the IC around the recording array.

### Optical and electrical stimuli

The light source was a custom-built solid state 488 nm laser (OptoTech, Melbourne, Australia) with light delivered through a 105 μm core silica optical fibre with 125 μm silica cladding, 3 mm PVC jacket and numerical aperture of 0.22 (Thorlabs, Newton, NJ). The fibre was connected to the laser using an FC connector and the distal end was bare fibre. The fibre was bonded to a 25 µm 90/10 Platinum/Iridium wire (insulation removed) with cyanoacrylate.

Stimulus waveforms were generated by an in-house purpose-built multichannel stimulator controlled by custom software implemented in Igor Pro (Wavemetrics, Portland, OR). Electrical current was delivered in Current Levels ‘CL’, where current in µA is given by: 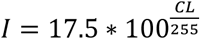. Light was presented as single pulses at 0-15 mW light intensities and 0.5-50 ms pulse widths. Irradiance was calibrated with a Fieldmaster power meter and LM10 power meter head (Coherent, Santa Clara, CA). Biphasic electrical stimulation (100 µs/phase with 50 µs interphase gap) was presented at sub- and suprathreshold levels (20-120% of threshold) with varying delays both during and after the optical pulses. Pulse train stimuli were presented in 5-240 Hz bursts to examine the rate-following efficiency of neurons to repetitive optical, electrical and combined stimuli. For acoustic IC recordings in normal hearing mice, acoustic stimulation was at 10-100 dB SPL, 4-32 kHz, 5 ms rise/fall and 100 ms duration.

### Data acquisition and analysis

Multiunit spike activity from the recording electrodes was amplified, filtered and digitized at a sampling rate of 33 kHz using a Cerebus data acquisition system (Blackrock Microsystems, Salt Lake City, UT). Multi-unit activity was processed using customised spike detection scripts in Igor Pro (Wavemetrics, Portland, OR) and is described in detail in (George et al., 2014). Briefly, a level of four times the root mean square for each recording channel was used to detect spikes. Spike counts were generated from a 5 – 40 ms window post stimulus, except for the pulse trains which used a 6-10 ms window. Spike counts across the array were used to display response images, with the depth of the IC recording array on the x-axis and the stimulus intensity on the y-axis. A spatial tuning curve (STC) image was generated for specific stimulation sites or acoustic frequencies. Threshold was determined for each condition, defined as the lowest stimulus intensity to elicit a normalised spike rate of 0.3. The recording site with the lowest threshold was defined as the best recording site.

The discrimination index (*d’*) represents the growth in neural response with increasing stimulus intensity at each recording site. At the best recording site, the value of d’ was cumulated across increasing stimulus levels above IC threshold. The widths of the STCs were measured at cumulative d’=1 and d’=2 for comparison between modalities.

Response in the IC to 300 ms duration bursts of stimuli presented at the cochlea was used to evaluate temporal fidelity. The maximum following rate was defined as the rate for which 80% of triggers had at least one spike in response. To avoid overlapping windows at high pulse rates, a fixed 6-10 ms post stimuli window was used to count spikes. Maximum following rates were determined with suprathreshold optical and subthreshold electrical stimulation at a level where d’ ≥ 2 level above threshold.

### Histology

Deeply anaesthetised mice were intracardially perfused with 0.9% (w/v) saline containing 0.1% (w/v) heparin sodium (ProVet, Melbourne, Australia) and 0.025% (w/v) sodium nitrite (Sigma-Aldrich, St Louis, MO), followed by neutral buffered formalin (NBF; Sigma-Aldrich, St Louis, MO). The temporal bones were extracted and the bulla was dissected away to expose the cochlea and semi-circular canals. The cochlea was post-fixed for a further 2 hours prior to rinsing with phosphate buffered saline (PBS) and decalcification in 0.12 M ethylenediaminetetraacetic acid (EDTA; Sigma-Aldrich, St Louis, MO) for 48 hours on an orbital shaker at 4 ^°^C. Decalcified cochleae were either frozen in optimal cutting temperature compound (OCT; ProSciTech, Thuringowa Central, Queensland, Australia) and sectioned at 12 µm through the mid-modiolar plane for -20 ^°^C storage or were microdissected for cochlear wholemount preparation.

Frozen sections were thawed at room temperature, incubated in PBS for 10 minutes, permeabilised in PBS containing 0.2% Triton X-100 (Sigma-Aldrich, St Louis, MO) and blocked in PBS containing 10% goat serum (Sigma-Aldrich, St Louis, MO) and 0.2% Triton X-100 and for 1-2 hours. A mouse monocloncal antibody to the C-terminus of ChR2 (#651180, Progen, Heidelberg, Germany) was applied to the sections for 2 hours, diluted 1:200 in blocking buffer. After washing the sections in PBS over 30 minutes, a secondary antibody was applied to the sections for 2 hours (AlexaFluor goat anti-mouse 594 diluted 1:400 in blocking buffer; Life Technologies, Carlsbad, CA). Sections were again washed in PBS over 30 minutes prior to the application of mounting media and coverslips.

Wholemounts were blocked in 0.1% Triton X-100 and 10% heat inactivated donkey serum (Sigma-Aldrich, St Louis, MO) in PBS for 1 hour. Rabbit polyclonal Myosin-VIIA antibody for hair cells (#25-6790; Proteus Biosciences, Ramona, CA) was diluted 1:200 in blocking solution and applied overnight at 4 ^°^C. Wholemounts were then treated with secondary antibody (Alexa 488 or Alexa Fluor 568 conjugated; Thermo Fisher Scientific, Scoresby, Victoria, Australia) diluted at 1:500 in PBS for 2 hours. The wholemounts were washed three times for 5 minutes in PBS and mounted in ProLong Diamond Antifade Mountant with DAPI (#P36965; Thermo Fisher Scientific, Scoresby, Victoria, Australia). All staining was visualized and imaged using the Zeiss fluorescence microscope and AxioVision software (Carl Zeiss, Oberkochen, Germany).

### In vitro electrophysiology

Cultures of dissociated spiral ganglion neurons were prepared from postnatal day 5 ChR2-H134R transgenic mice (n=12) as described by (Hart et al., 2020). Electrophysiology recordings were carried out following previously established protocols (Hart et al., 2020). Briefly, glass micropipettes were pulled from thick-walled borosilicate (GC150F; SDR Scientific, Sydney, Australia) using a Sutter P87 Flaming Brown Micropipette Puller (Sutter Instruments, Novato, CA) for 5 – 10 MΩ resistance. The intracellular solution consisted of (in mM) K-glu 115, KCl 7, HEPES 10, EGTA 0.05. Sucrose was used for osmolarity (305-310 mOsm) and 2.5 M KOH to adjust pH to 7.1 – 7.3. Extracellular solution consisted of (in mM) NaCl 137, KCl 5, HEPES 10, D-Glucose 10, CaCl_2_, MgCl_2_. Sucrose was used to adjust osmolarity (305-310 mOsm) and 10 M NaOH to adjust pH to 7.3 – 7.4. All reagents were purchased from Sigma-Aldrich, St Louis, MO. A junction potential of 12.8 mV was subtracted from all recorded potentials.

Recording was carried out using a Multiclamp 700B digitizer (Molecular Devices, San Jose, CA) and Digidata 1440A amplifier (Molecular Devices, San Jose, CA) with a CV-7B head stage. An OptoPatcher (A-M Systems, Carlsborg, WA) was used to align the fibre through the patch pipette. Recordings were made using Axograph software. Laser settings were controlled by a custom software interface connected by USB, and pulses were externally triggered by a Multiclamp 700B TTL output. The light source was the same as for the *in vivo* studies. A triple pulse protocol was applied, in which a hybrid optical/electrical pulse was delivered, followed by isolated 0.3 ms electrical and 1 ms optical pulses at the same stimulus level. Presentations where the isolated stimuli evoked action potentials were excluded.

## Results

### Animal model

To examine the spectral resolution and temporal fidelity of optical, electrical and hybrid stimulation of the auditory system, we used transgenic mice that expressed the ChR2-H134R-EYFP fusion protein via the parvalbumin promoter. Expression of ChR2-H134R was examined via a ChR2 antibody and confirmed to mirror the localisation of EYFP (enhanced yellow fluorescent protein) in spiral ganglion neuron cell bodies, peripheral fibres and central fibres. The strongest fluorescence localised to the membrane of spiral ganglion neurons (**Figure 1(a)**). Additionally, ChR2-H134R was present in inner hair cells with weaker expression in the outer hair cells of adult transgenic mice (**Figure 1(a)**). This expression pattern was evident throughout the cochlea, from the basal turn to the apical turn. The mice were acutely deafened with neomycin to reduce the possibility of hair cell-mediated auditory responses. The efficacy of the acute neomycin deafening procedure was verified histologically in a subset of mice which showed complete or near complete loss of inner and outer hair cells (n=2; **Figure 1(b-d)**).

**Figure 1.**
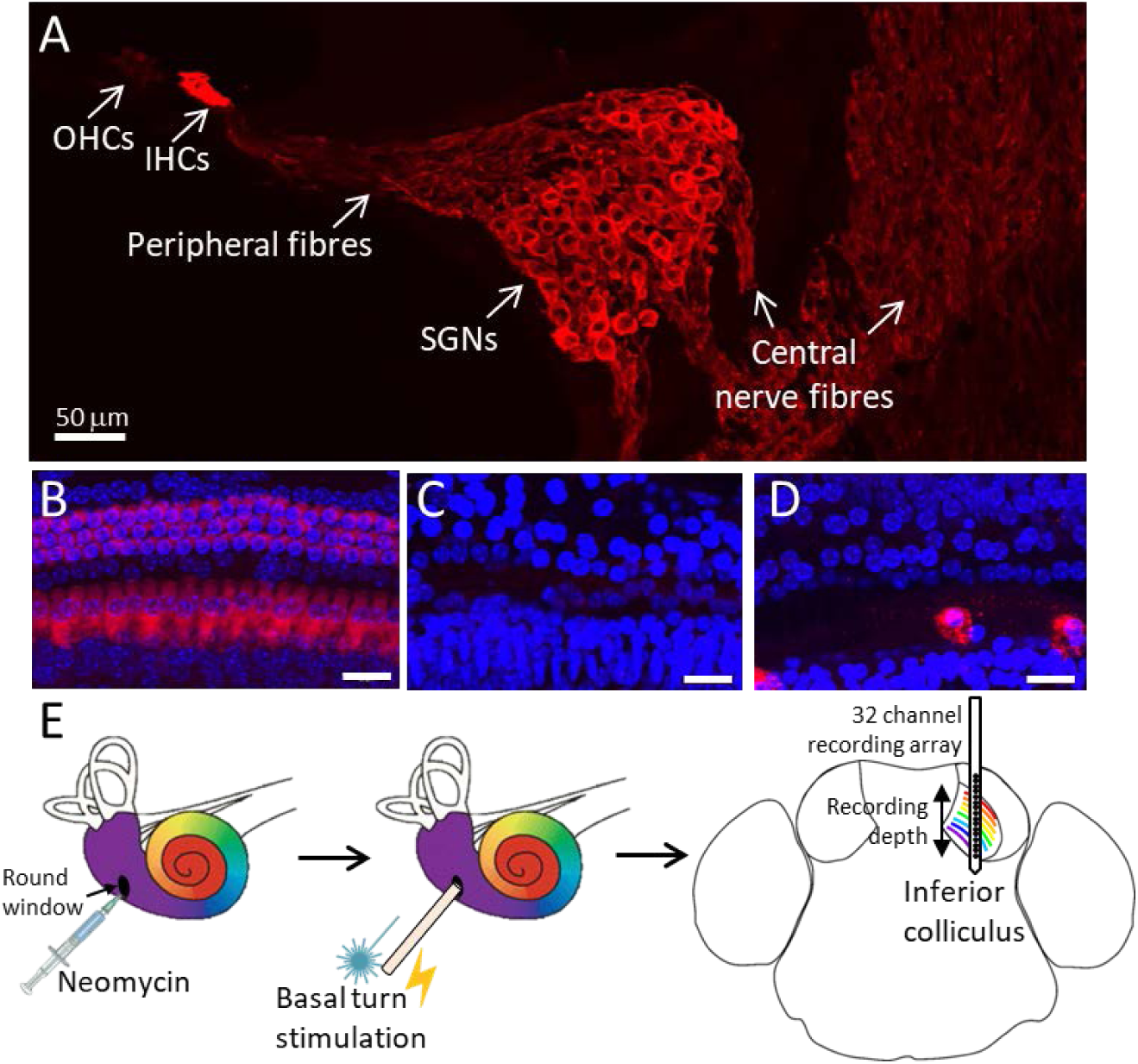
Optogenetic mouse model with acute hearing loss. **(A)** Cochlear section from ChR2-H134R transgenic mouse stained with ChR2 antibody (red) showing expression in spiral ganglion neuron (SGN) cell bodies, peripheral fibres and hair cells. **(B)** Representative image of the sensory epithelium in the 16 kHz region of a normal hearing mouse cochlea (red = Myosin VIIA staining hair cells, blue = DAPI). Two weeks after exposure to neomycin there was complete loss of inner and outer hair cells **(C)** or sporadic survival of inner hair cells (red) **(D). (E)** Experimental protocol: Transgenic mice were acutely deafened by infusion of 100 mg/mL neomycin solution through the cochlear round window. An optical fibre with an attached platinum wire was inserted 1-2 mm into the round window. During stimulation, multiunit responses were recorded from the inferior colliculus of the auditory midbrain via a 32-channel recording array. Scale bars for B-D are 20 µm. IHCs - inner hair cells, OHCs – outer hair cells, SGNs – spiral ganglion neurons.

### Activation thresholds

Optical pulses (1 ms), biphasic electrical pulses (250 µs) or hybrid stimulation was delivered to the cochlear spiral ganglion neurons of acutely deafened transgenic mice using an optical fibre and platinum wire inserted into the round window of the cochlea. During stimulation, neural recordings were made from the IC of the auditory midbrain using a 32 channel recording array (**Figure 1(e)**). Threshold power levels were defined for optical and electrical pulses, respectively, as the lowest stimulus power level required to elicit a normalised spike rate of at least 0.3 (that is a 30% increase between spontaneous and maximum firing rates). Hybrid pulses were then presented at varying levels above and below threshold. The start of the electrical pulse was delayed with respect to the start of the optical pulse by a variable time t_d_ so that when t_d_ = 750 µs both the optical and electrical pulses ended simultaneously. The delays tested were 0 µs (i.e. no delay), 750 µs, 1750 µs, 2750 µs and 3750 µs. **Figure 2** shows an example raster plot indicating when a neural response was detected in response to 10 repeats for sub- and suprathreshold stimulation levels. It can be seen that the sub-threshold optical and sub-threshold electrical stimuli applied individually did not evoke responses. By contrast, the application of these stimuli in combination elicited neural activity in the IC when t_d_ ≥ 750 µs. The reduction in electrical threshold that could be achieved with a sub-threshold optical pulse is shown in **Figure 3** as determined from n=9 cochleae for different delays. When the start of the electrical pulse coincided with the start of the optical pulse (no delay; t_d_ = 0), there was minimal effect on thresholds. When the start of the electrical pulse was delayed after the start of the optical pulse, neural responses were detected with subthreshold levels of both optical power and electrical power. Significant reductions in electrical threshold were measured for delays of up to 3.75 ms where the electrical pulse occurred after the end of the optical pulse (p<0.05; ANOVA, n=9) (**Figure 3**).

**Figure 2.**
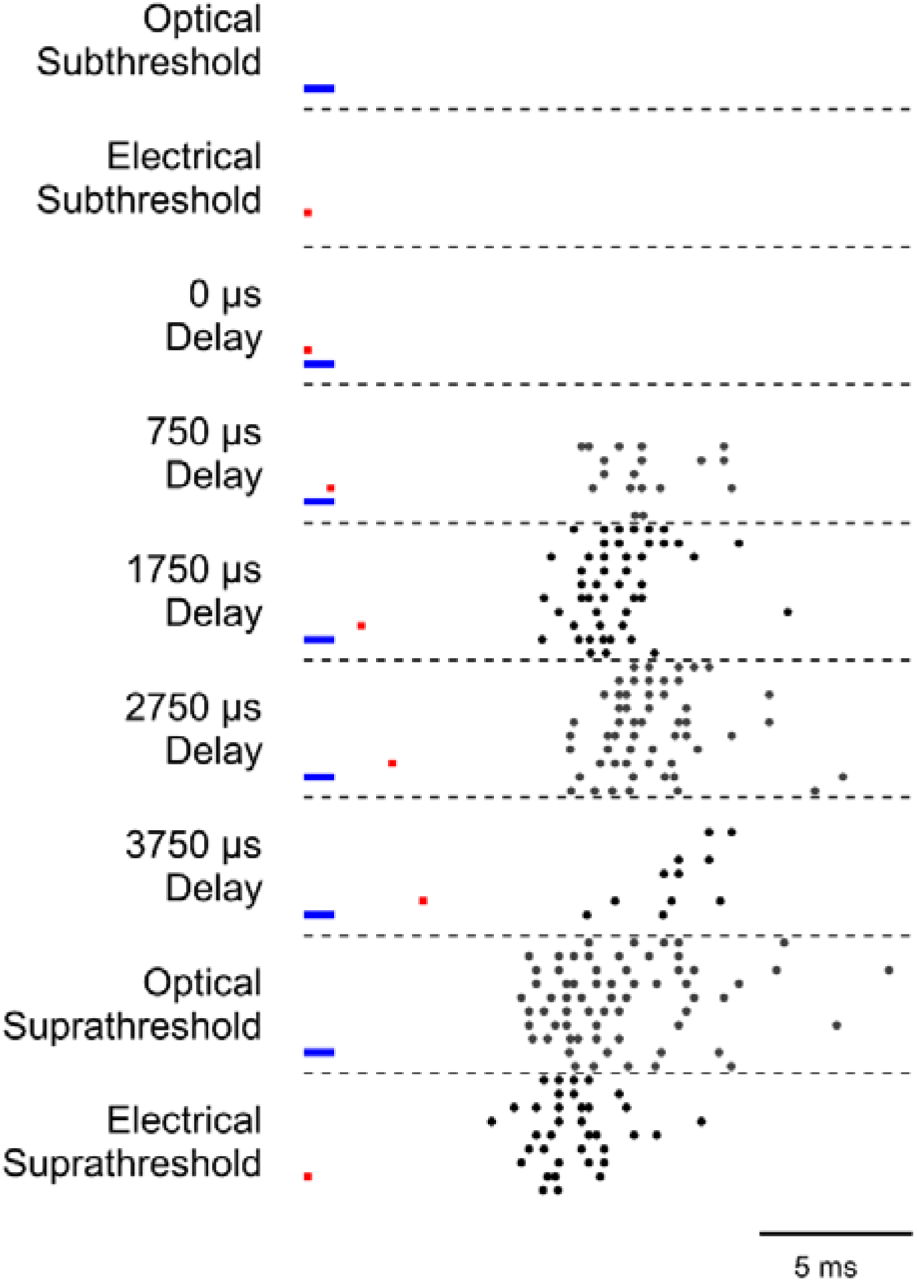
Activation of the auditory pathway by hybrid sub-threshold optical and sub-threshold electrical stimuli. Raster plot showing the response to subthreshold for optical alone, electrical alone and hybrid stimulation at the same stimulus levels as the individual stimuli with t_d_ = 0, 750 µs, 1750 µs, 2750 µs and 3750 µs. In this example, the electrical stimulus (70 Current Levels; CL) was 64% of threshold (95 CL) and the optical stimulus (25 µW) was 70% of threshold (35 µW). Pulse timing is shown with blue (optical) and red (electrical) lines. When combining subthreshold stimuli, neural responses were only evoked when the electrical stimuli were delayed at least t_d_ = 750 µs to end simultaneously with the optical pulse and a maximum response was observed for t_d_ = 1750 and 2750 µs. For comparison, responses to individual suprathreshold stimuli (50 µW and 100 CL) are also shown.

**Figure 3.**
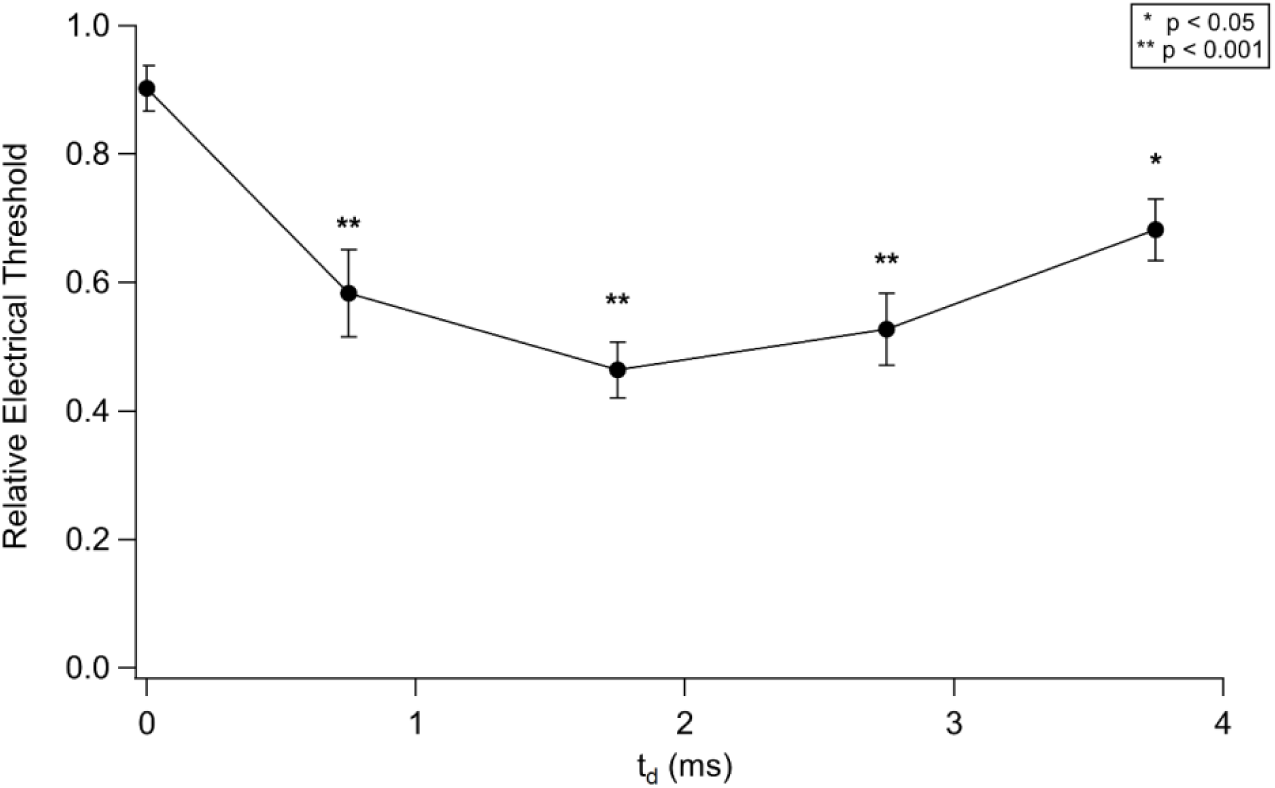
The timing of the electrical pulse affects the reduction in the electrical threshold observed for hybrid stimulation. The ratio of electrical threshold with no optical pulse compared to electrical threshold with a sub-threshold optical pulse at delays of 0 to 3.75 ms is shown. Facilitation of subthreshold optical stimulation occurred when the subthreshold electrical stimulus delay was 0.75-3.75 ms. Compared to no delay (t_d_ = 0 µs), a significant reduction in relative electrical threshold was observed when t_d_ ≥ 750 µs (p < 0.05 ANOVA). Results from n=9 cochleae, errors bars show standard error.

In addition, electrophysiology recordings were performed on cultured spiral ganglion neurons from ChR2-H134R transgenic mice to examine spike shape and the post-stimulus facilitation period in single cells, using 1 ms optical pulses and 300 μs monophasic electrical pulses. Threshold stimulus was defined as the power level where action potentials were evoked for at least half of the presented stimuli. For individually presented single pulses, there was a dramatic difference in the response of the neurons to subthreshold optical stimuli and subthreshold electrical stimuli. On average, spiral ganglion neurons responded to subthreshold electrical stimuli with sharp depolarisation followed by a rapid decay immediately after the cessation of electrical stimulus. In contrast, the response to subthreshold optical stimuli was a slower, smaller depolarisation which continued after the cessation of optical stimulus and returned to the resting membrane potential over 30-50 ms (**Figure 4(a)**). Hybrid stimuli were then presented (electrical at 30-80% of threshold and optical at 80-100% of single pulse threshold) with a variable delay. When the electrical pulse preceded the optical pulse (*t*_*d*_ <0), firing probability was less than 40%. Where the electrical pulse lagged the optical pulse (*t*_*d*_ >0), the firing probability increased to a maximum (83%) for delays of 9 to 13 ms. Firing probability then decayed exponentially from the peak, falling below 50% firing probability when the electrical pulse was applied more than 30 ms after the start of the optical stimulus. Action potential facilitation (defined as 30 – 40% firing probability) persisted for up to 60 ms after the start of the optical stimulus (**Figure 4(b)**).

**Figure 4.**
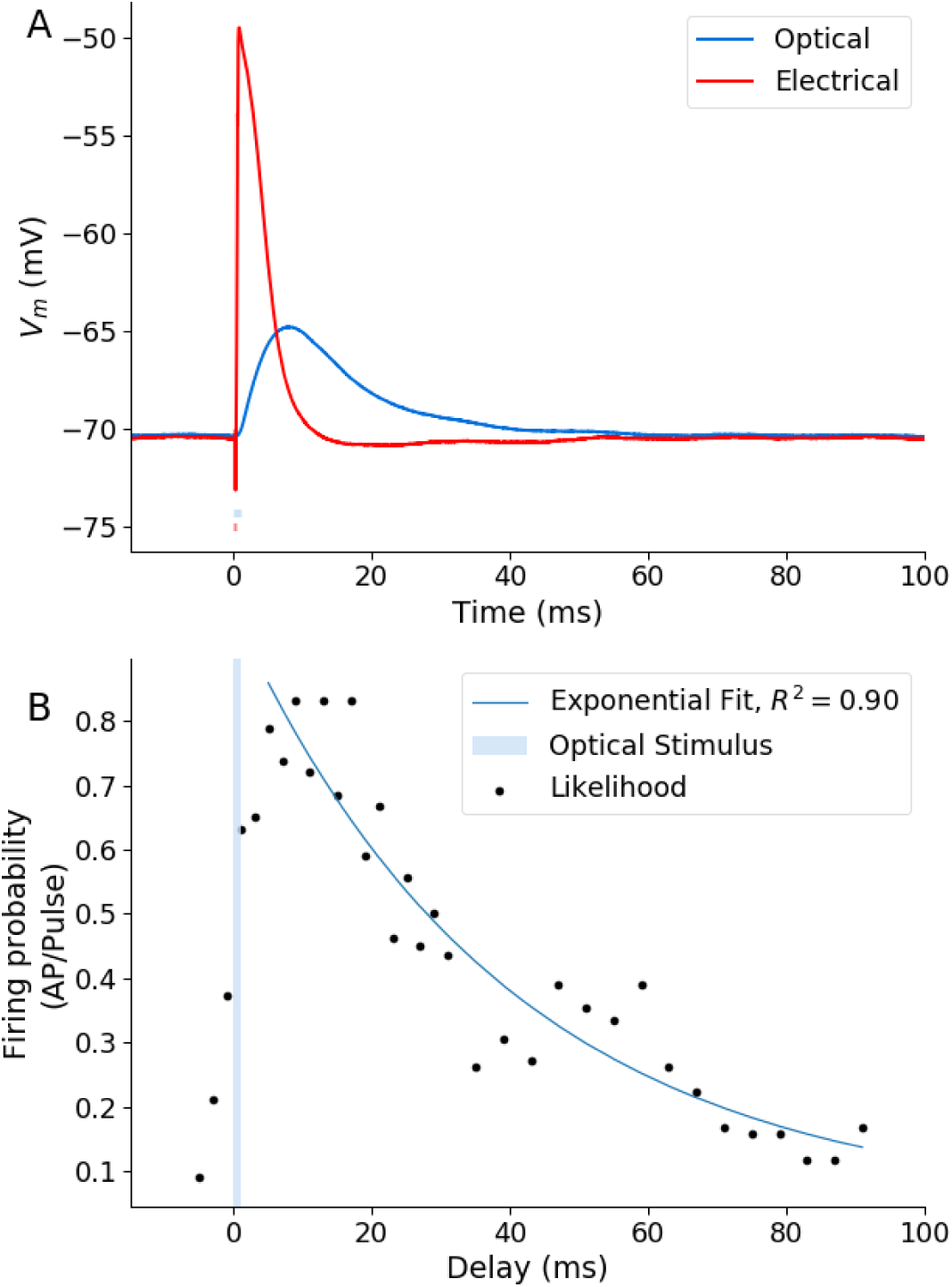
Post-stimulus facilitation during hybrid stimulation reflects the change in membrane potential of a subthreshold optical pulse. **(A)** The average response of single spiral ganglion neurons to separate subthreshold electrical (red; 300 µs) and subthreshold optical (blue; 1 ms) stimuli, with the pulses indicated below the response curves. The response to electrical stimuli showed a rapid depolarisation followed by a rapid decay after the cessation of stimulus. The response to the optical stimulus was a slow increase in membrane potential, peaking at approximately 8 ms after the cessation of stimulus, followed by a slow decay. **(B)** The probability of firing an action potential (AP) for a 1 ms subthreshold optical pulse (blue shaded region) and a 300 µs subthreshold electrical pulse applied before or after the start of the optical pulse with varying delays (n=4 cells, 852 presented stimuli, electrical at 30-80% threshold, optical at 80-100% threshold excluding the cases where the 100% optical-only stimulus produced an action potential). The dashed line is an exponential decay curve fitted to the individual points where the delay t_d_ ≥ 5 (y =0.92 · exp(−0.025x) +0.04, R^2^ =0.9). This suggests a decay time constant of 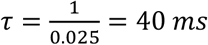.

### Entrainment rates

To evaluate the temporal fidelity of hybrid stimulation compared to optical alone we assessed the response in the IC to 300 ms bursts of stimuli presented at the cochlea. For this study, hybrid stimulation comprised suprathreshold optical stimulation and varying levels of subthreshold electrical stimulation in order to make direct rate-following comparisons to suprathreshold optical-only stimulation. **Figure 5** shows an example raster plot from suprathreshold optical only and hybrid stimulation (suprathreshold optical at 12.5 mW, subthreshold electrical at 140 Current Levels; CL, which was 83% of threshold). At 40 pps, both stimuli had robust entrainment, but at 80 pps only the hybrid stimuli showed entrainment. At the maximum rate tested (240 pps), there were very few spikes recorded after the initial onset from the optical-only stimuli. By contrast, at 240 pps the hybrid stimulation was able to evoke responses over the whole 300 ms window.

**Figure 5.**
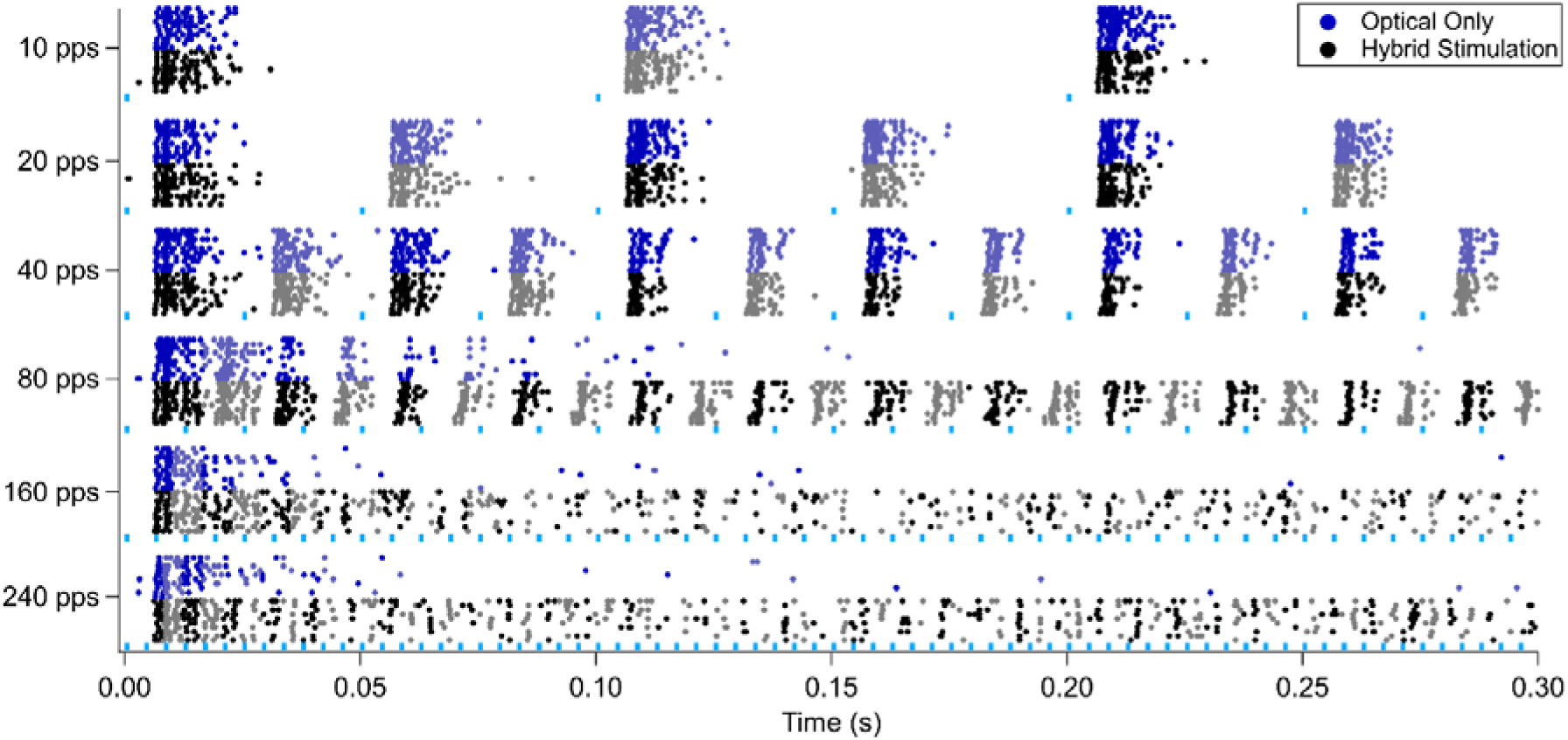
Improved entrainment achieved with hybrid stimulation compared to optical only stimulation at higher stimulation rates. Raster plot comparing responses to suprathreshold optical-only stimulation and hybrid stimulation (suprathreshold optical, subthreshold electrical) presented at rates between 10 and 240 pulses per second (pps). Laser pulses are shown in light blue bars underneath each pulse rate.

To better quantify the stimulation, the maximum following rate was found where 80% of triggers had at least one spike in response. To avoid overlapping windows at high pulse rates, a fixed 6-10 ms post stimuli window was used to count spikes. Maximum following rates were determined with suprathreshold optical and subthreshold electrical stimulation (typically at a level where the discrimination index (d’) ≥ 2 level above threshold, where d’ is a measure of sensitivity or discriminability derived from signal detection theory).

An example of the percent following for suprathreshold optical stimulation alone (9.75 mW) and hybrid stimulation (suprathreshold optical with increasing levels of electrical from 40% of threshold to 83% threshold) is shown in **Figure 6(a)**. As the electrical current was increased, the percent following increased and the maximum following rate shifted from 51 Hz to 122 Hz in this example. **Figure 6(b)** shows the average increase in following rates when using hybrid stimulation compared to optical only (n=4). For these stimuli, electrical was always sub-threshold and optical was at a level to give d’ ≥ 2 above threshold. With hybrid stimulation a significantly greater maximum following rate of 111 Hz was achieved, compared to just 46 Hz for optical alone (p<0.05; paired T-test, n=4).

**Figure 6.**
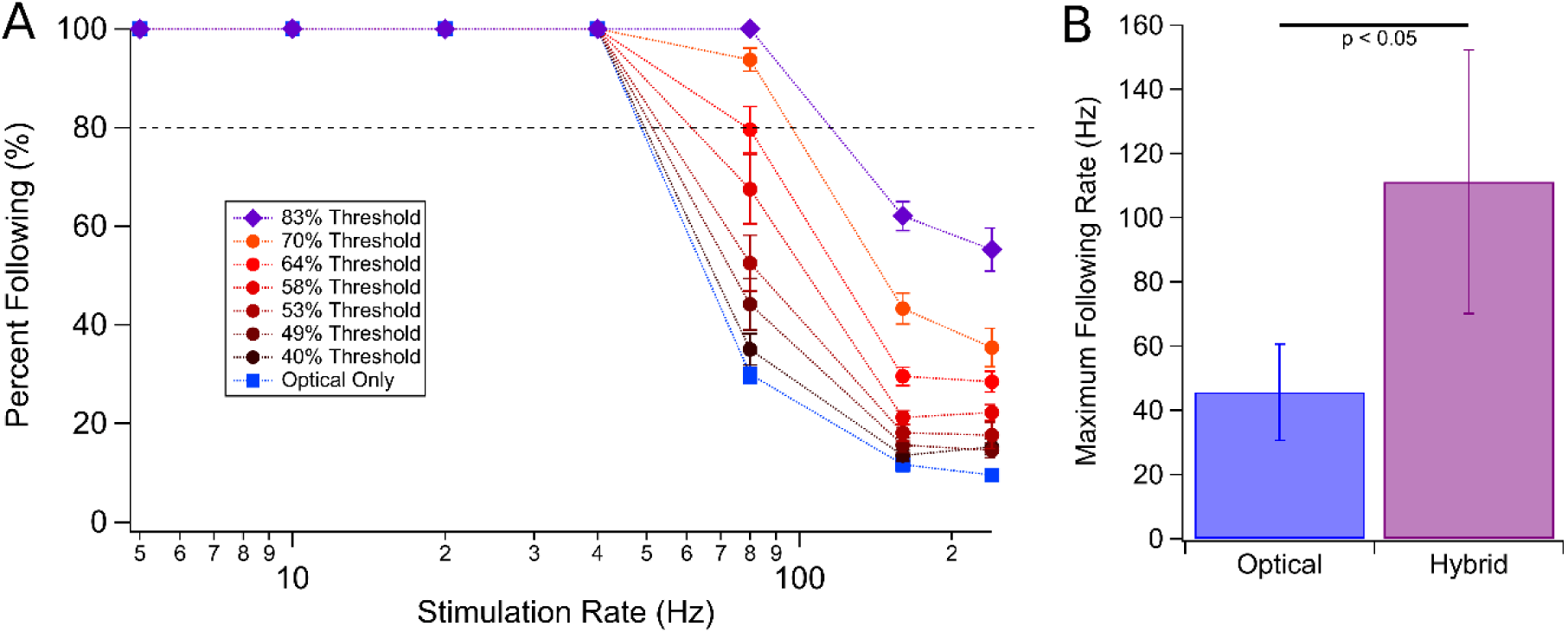
The average following rates are greater when using hybrid stimulation compared to using optical stimulation alone. **(A)** Percent following for suprathreshold optical stimulation alone and hybrid stimulation comprising suprathreshold optical pulses (9.75 mW) with increasing levels of electrical current from 40% of threshold to 83% of threshold (where threshold was 150 CL) at rates of 5 – 240 Hz. In this example, at the best performing level of hybrid stimulation with subthreshold electrical, 55% of stimuli had at least one spike in response at 240 Hz (purple diamonds), while only 9.6% of stimuli had a response with optical only (blue squares). **(B)** Maximum following rate for optical and hybrid stimulation. The hybrid stimulation has a significantly higher maximum following rate (111 Hz) compared to optical alone (46 Hz; p<0.05, n=4 cochlea, paired T-Test) Error bars show standard error.

### Spread of activation

We next wanted to evaluate the efficacy of hybrid stimulation for improving the spectral resolution of activation compared to electrical only stimulation. In this case hybrid stimulation consisted of a fixed level of subthreshold optical stimulation and varying levels of electrical stimulation from sub- to suprathreshold in order to make direct comparisons of excitation width to varying levels of electrical-only stimulation. An image of the spatial extent of evoked multi-unit activity across the recording array in the IC of a normal hearing ChR2-H134R transgenic mouse was generated in response to acoustic stimuli (n=1). Similar images were generated of the IC response in acutely deafened ChR2-H134R transgenic mice to electrical, optical and hybrid stimuli (n=9), with example responses shown in **Figure 7** to the four stimulation types. In each image the firing rate is indicated by colour, where black indicates spontaneous neural firing rate and yellow indicates the maximum firing rate. Optical stimulation pulses were 1 ms, biphasic electrical pulses were 250 µs and hybrid stimulation used t_d_ = 750 µs so the optical and electrical pulses ended together. **Figure 7(a)** shows a sharply tuned STC to a 28 kHz pure tone acoustic stimulus for a normal hearing ChR2-H134R transgenic mouse. **Figure 7(b-d)** present IC response images for basal turn electrical, hybrid and optical stimulation in an acutely deafened mouse, where the optical fibre and platinum wire were inserted 2 mm through the round window membrane. All modalities resulted in the lowest thresholds at the more ventral recording electrodes, which is indicative of activation at the base of the cochlea. Unlike acoustic stimulation, the extent of multiunit activity for electrical only and optical only stimulation increased with increasing stimulus intensity above threshold. Hybrid stimulation resulted in a narrow band of activation at the ventral recording sites but also high threshold activation at the more apical recording sites. Optical responses were not detected in mice that did not express ChR2-H134R (data not shown).

**Figure 7.**
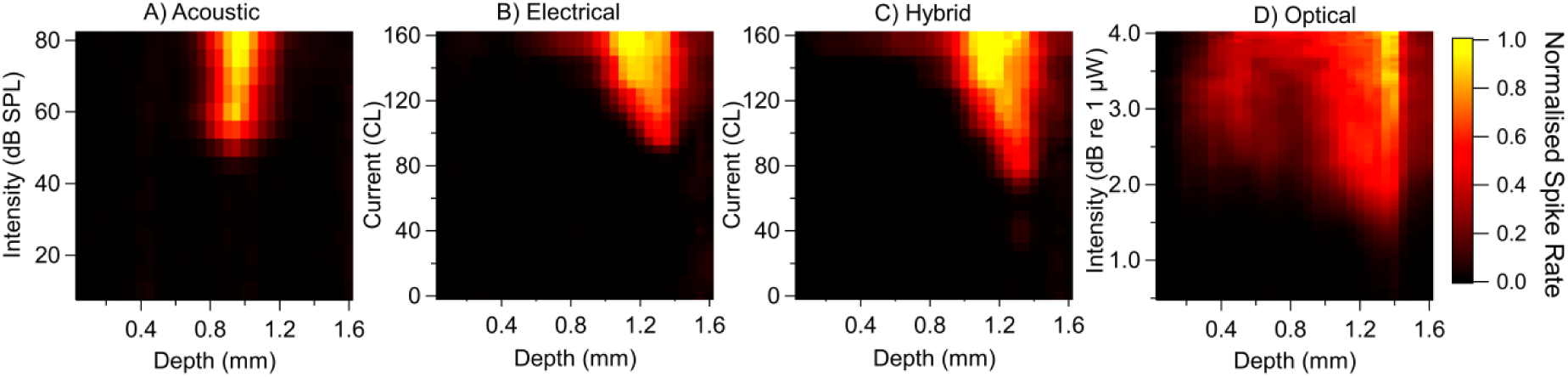
Hybrid stimulation significantly reduces spread of activation compared to electrical or optical-only stimulation. Example of response images illustrating the spatial extent and rate of multi-unit activity across the recording sites of the recording array in the IC (measured as recording depth) in response to: **(A)** 28 kHz acoustic pure tones; **(B)** 100 µs/phase electrical stimuli **(C)** Hybrid stimulation at highest optical level that remained subthreshold (25 μW) with the electrical pulse delayed to finish with the optical pulse (td = 750 µs) **(D)** 1 ms duration optical-only stimulation. Each point on the response images was normalized between the spontaneous activity rate (black) and maximum response (yellow). Optical power was converted from mW to a linear scale (dB referenced against 1 uW) for comparison to the other modalities. CL = current levels. Response images from B-D were recorded from the same animal.

The spread of activation was compared for acoustic, optical, electrical and hybrid stimulation applied to the base of the cochlea. The width of excitation in the STC was assessed at intensities of levels of d’ = 1 and d’ = 2 above threshold to control for differences in linearity of stimuli as illustrated in **Figure 8(a)**. Electrical and optical stimulation resulted in activation in the IC that was spatially broader than acoustic stimulation. There was no significant difference between optical only and electrical only stimulation. In contrast, the width of activation in the IC was significantly reduced for hybrid stimulation compared to with electrical stimulation or optical-only stimulation at d’=2 (p<0.01, ANOVA, n=9; **Figure 8(b)**).

**Figure 8:**
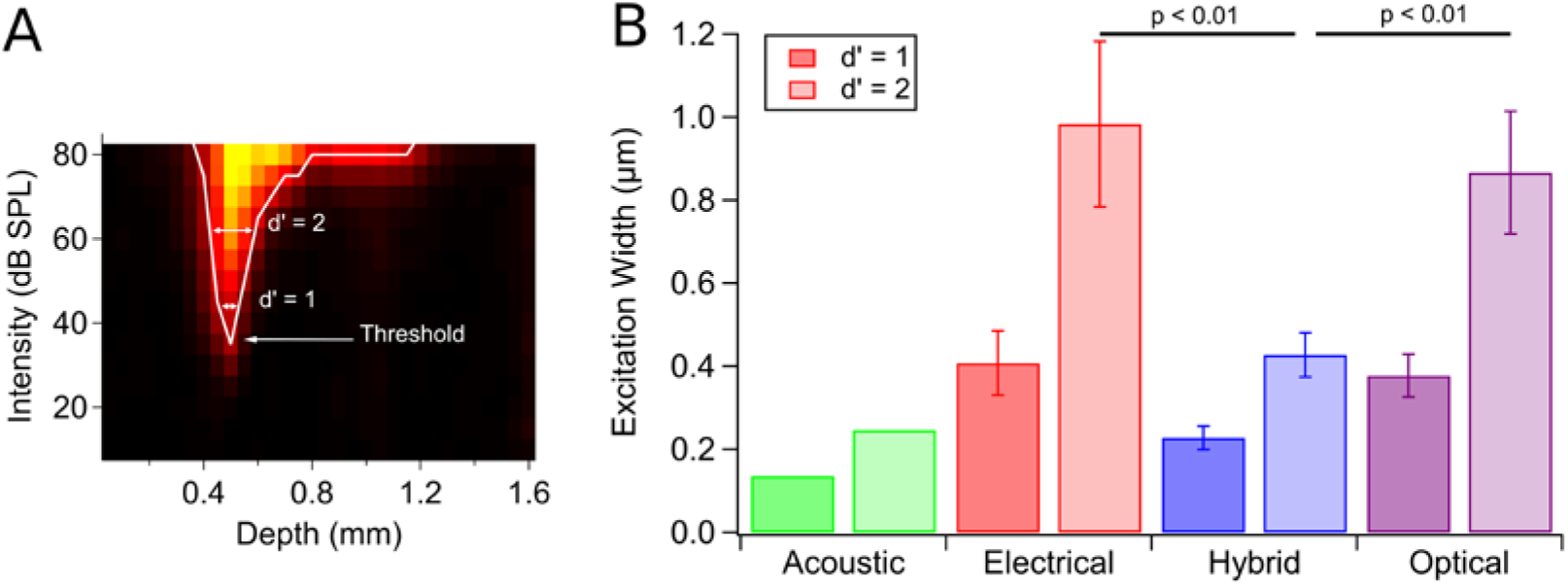
Hybrid stimulation significantly reduces spread of activation compared to electrical or optical-only stimulation at d’=2 above threshold. **(A)** An example acoustic IC response image illustrating the spatial tuning curve (STC) generated by connecting the stimulus levels that elicited 0.3 normalized responses on each IC recording site (white line). The tip of the STC is the best threshold. Excitation widths were measured at d’=1 and d’=2 above threshold. **(B)** Spread of activation for different stimulation modalities at d’ = 1 and d’=2. Hybrid stimulation had significantly narrower activation in the IC when compared to optical and electrical alone at the d’ = 2 level (p<0.01, RM ANOVA, n=9) Error bars show standard error.

## Discussion

In this study we demonstrated that hybrid stimulation in the cochlea, i.e. the combination of optogenetic and electrical stimulation when one or both modalities were at sub-threshold levels, significantly improved the spectral resolution *and* temporal kinetics (maximum following rate) of neural activation compared to electrical-only or optical-only stimulation, respectively. The combination of sub-threshold electrical and sub-threshold optical stimuli resulted in activation of the auditory pathway *in vivo* and within individual spiral ganglion neurons *in vitro*, thus reducing the optical radiance required for neural activation compared to optical-only stimulation. This facilitation was most apparent when the electrical stimulus was delayed with respect to the start of the optical pulse and could extend beyond the completion of the optical pulse, indicating post stimulus facilitation with optical stimuli. These results suggest that hybrid stimulation can be used to improve temporal fidelity in neurons expressing opsin ion channels while improving spectral resolution.

### Hybrid stimulation

Our data shows that sub-threshold optical stimulation and sub-threshold electrical stimulation can be combined to result in action potentials in auditory neurons *in vitro* using shorter stimuli than our previous work on auditory neurons (Hart et al., 2020), and activation of the auditory pathway *in vivo* which has not been reported before. Hybrid stimuli have been examined previously for the rodent sciatic nerve where sub-threshold electrical stimuli were combined with either sub-threshold infrared neural stimulation (INS) or optogenetic-mediated blue light to result in muscle activation (Duke et al., 2009, Kapur et al., 2016), reducing the radiant energy required for neural activation (Duke et al., 2012b). For INS, the hybrid stimulation effect was maximal when the pulses were applied simultaneously or when the pulses ended at the same time (Duke et al., 2009, Duke et al., 2012b). This contrasts to the timing for maximum probability of hybrid activation in the cochlea of our optogenetic mice, where peak multiunit activity in the IC occurred when a delay of 2 ms was applied between the start of the optical pulse and the start of the subsequent electrical pulse *in vivo*. This difference reflects the different mechanisms of activation of INS and optogenetic stimulation, whereby the closing kinetics of the ChR2-H134R ion channel (τ_off_ ∼ 18 ms) influences the period of excitability of the neuron.

We used patch clamp electrophysiology to understand hybrid stimulation mechanisms at the cellular level. Our data showed the rate of repolarisation following the optical stimulus was reduced relative to electrical-only stimulation due to persistent photocurrents beyond the optical pulse. Post-stimulus facilitation in single auditory neurons extended to delays of 9-13 ms between the optical and electrical stimuli *in vitro*, which is greater than the facilitation observed *in vivo*. Dynamics and firing rates are generally slower *in vitro*, as found previously for electrical stimulation (Wright et al., 2016). Despite these differences, the observed facilitation both *in vitro* and *in vivo* suggests that cultured SGNs are a good *in vitro* model for hybrid optical stimulation. Post-stimulus facilitation, or summation, is also seen with electrical-only stimulation, although the facilitation was much less persistent for electrical-only stimulation compared to hybrid stimulation (Hart et al., 2020). In the cat, facilitation is described for a sub-threshold electrical pulse followed by another electrical pulse with inter pulse intervals of less than 1 ms (Cartee et al., 2000) and with high rate stimulation in the guinea pig, where significantly reduced thresholds were observed, indicative of interactions between pulses (Heffer et al., 2010).

A key benefit of combining optical stimulation with electrical stimulation is the reduced optical radiance required to activate neurons compared to optical-only stimulation. This is important when considering the overall safety of optical stimulation in the cochlea and potential issues with phototoxicity and thermal trauma. Phototoxicity is dependent on the wavelength of the light, with blue light considered more harmful than red. While the phototoxicity of both blue and red light has been found to be low-risk for brain tissue at stimulation levels in excess of those considered here (Senova et al., 2017), the effect of life-long optical stimulation is unknown. Heating considerations in the confined space of the cochlea will be dependent on the ultimate design of an optical cochlear implant. For a micro-LED-based system, a temperature increase of less than 0.5 ^°^C was observed in brain tissue at rates of 500 Hz using light intensities far in excess of those required to activate opsin-expressing neurons (McAlinden et al., 2013). Thresholds for optogenetics-based stimulation in the auditory system vary depending on the optical fibre diameter, the opsin and its expression levels. Therefore, there is a wide range of radiant exposure levels reported for neural activation (reviewed in (Richardson et al., 2020)). Hybrid stimulation will increase overall safety of optical stimulation by lowering the required radiant flux due to reduced thresholds of activation and the extended period of excitability beyond the optical pulse during which electrical stimuli can be used to activate neurons.

### Entrainment rates

One limitation of optogenetics-based optical stimulation methods is low temporal fidelity which is dictated by the opening and closing kinetics and post-stimulus refraction periods of each opsin. The maximal stimulation rate is dependent on neural subtype but is approximately 70 Hz for wild-type ChR2 *in vivo* (Hernandez et al., 2014). This is problematic for the auditory system in which complex-tone pitch is encoded not only by place of excitation but also by the stimulus repetition rate. Cochlear implant recipients can typically discriminate up to 300 Hz (reviewed in (Zeng, 2004)), but no benefit in speech perception was observed beyond 500 Hz (Shannon et al., 2011). One solution is to use opsins with fast on-off channel kinetics such as Chronos which significantly improved spike probability and increased stimulation rates in the cochlea, with an average 50% spike probability at approximately 150 Hz and some neurons following the optical stimulus up to several hundred Hz (Mager et al., 2018, Keppeler et al., 2018, Duarte et al., 2018). As an alternative solution, the results presented here indicate that hybrid stimulation, whereby optical pulses are used to prime the spiral ganglion neurons and sub or suprathreshold electrical stimuli are used to activate the neurons, can boost the maximum following rate of auditory neurons expressing the ChR2-H134R opsin. Our data indicated a 2.4-fold increase in maximum following rate, a key measure of temporal fidelity, while lowering the radiance thresholds required for neuronal activation. If the effect of hybrid stimulation on entrainment rate can be extrapolated to other opsins, the number of candidate opsins that could be used for activation of the auditory system at rates up to 300-500 Hz can be expanded beyond Chronos alone. For example, the red-shifted ChR2 ET/TC also has high photocurrents and τ_off_ ∼ 8 ms (Berndt et al., 2011).

### Spread of activation

We did not observe any difference in the spread of activation between optical-only and electrical-only stimulation in this study. This contrasts with other studies which have shown statistically narrower STCs for optical-only stimulation compared to monopolar stimulation and bipolar stimulation (Dieter et al., 2019, Richter et al., 2011). In the gerbil, the angle of the fibre placement at the round window membrane was found to affect STCs, which was suggested to be a result of some angles causing light to penetrate through to more apical cochlear tissue (Dieter et al., 2019). Our findings are in agreement with a Monte Carlo ray tracing simulation which suggests that high intensity irradiance at the basal turn of the cochlea using a non-optimised fibre placement results in activation of apical neurons (Wrobel et al., 2018). In our study, the angle of fibre placement did not influence the STC, which may be a reflection of the difference in the animal model or in the light delivery method. In the gerbil model, opsin expression in neurons was achieved by viral-based gene therapy in which approximately 30% of neurons were transfected to varying degrees throughout the cochlea (Wrobel et al., 2018). In our heterozygous transgenic mouse model, all spiral ganglion neurons express ChR2-H134 with expression levels dictated by the parvalbumin promoter. Expression levels have been suggested to influence photosensitivity (Meng et al., 2019) which would affect activation thresholds and therefore spread of activation.

Suprathreshold optical stimulation in the basal cochlear turn resulted in low threshold activation of the layer of the IC corresponding to the basal region of the cochlea, but also high threshold activation corresponding to more apical areas, suggestive of penetration or scattering of the light beyond the basal site of stimulation. When we combined optical and electrical modalities, we observed a significant reduction in the spread of activation compared to optical-only stimulation at d’ values of 2 above threshold. This may be explained by the reduced strength of the optical radiance used during hybrid stimulation which would result in lower penetration of the light to the more apical regions. With the electrical stimulating level also being very low during hybrid stimulation, the overall result is a reduced spread of activation. This effect cannot be achieved with electrical-only stimulation as a certain number of neurons need to be activated for the sound to be perceived. Alternatively, it is possible that optical activation of the neurons is occurring centrally as well as at the peripheral fibres. The ChR2-H134 ion channel was expressed in the peripheral and central fibres and the cell body of spiral ganglion neurons. Thus an optical fibre positioned in the basal turn could activate the passing central fibres of spiral ganglion neurons from more apical cochlear regions, impacting the excitation pattern observed in the IC. This is assumed to occur for electrical stimulation as well (Cartee et al., 2000), in contrast to acoustic stimulation which occurs at the synaptic membrane.

The relative placement of the optical emitter and electrode *in vivo* could impact on the degree of interaction between the optical and electrical stimuli. In this study, the platinum wire was glued to the optical fibre so that they were in the same position in the cochlea. However, the optical fibre is orientation dependent, whereas the platinum wire is not. As the optical fibre is inserted through the round window membrane, the light path is suboptimal which would impact on the recruitment of auditory nerve fibres, as found previously (Duke et al., 2012a, Dieter et al., 2019, Wrobel et al., 2018). The current development of high density optical cochlear implants is based on LED chips or thin-film micro-LEDs in which the light is directed toward the cochlear modiolus where the spiral ganglion cell bodies are situated (Klein et al., 2018), with modelling suggesting that more optimised light sources such as these can have a significant impact on spectral resolution (Wrobel et al., 2018). In a device such as this, optimum hybrid activation will require the electrodes to be positioned close to each LED in alternating positions.

## Conclusions

Electrical stimulation is used by contemporary cochlear implants to activate neurons with high temporal fidelity, power efficiency and safety. However, it is almost impossible to achieve high spectro-temporal control because responsive regions overlap spatially and this in turn results in poor pitch discrimination of complex tones. Optical stimulation has the potential to deliver high spatial precision, but many optogenetic ion channels have poor temporal kinetics, require high optical radiance and presently have an unknown long-term safety record. Our results suggest that hybrid stimulation has improved temporal precision compared to optical stimulation alone, and greater spectral resolution than electrical stimulation alone. Therefore, this approach has the potential to increase the number of perceptually independent stimulating channels through reduced channel interactions, while reducing the overall light exposure levels and the associated risks of phototoxicity and thermal trauma.

## Acknowledgements

The authors acknowledge the technical support provided by Trung Nguyen, Brianna Flynn, Ella Trang, Caitlin Singleton, James Firth and Louise Rooney. Transgenic mice used in this study were generously donated by Prof Heather Young (University of Melbourne) and Prof Steven Petrou (The Florey Institute of Neuroscience and Mental Health). Viral vectors were provided by Prof Karl Deisseroth.

This work is supported by Action on Hearing Loss International Project Grant G89 and the ARC Training Centre in Biodevices (IC140100023). William Hart is supported by an Australian Government Research Training Program (RTP) Scholarship. The Bionics Institute acknowledges the support it receives from the Victorian Government through its Operational Infrastructure Support Program.

## Competing interests

The authors declare that a provisional patent has been filed in this area.

## Author contributions

Conception and study design: RR, AW, JF, KN, PS

IC recordings, data analysis: AT, RR, JF

In vitro electrophysiology recordings and analysis: WH, KN

Histology: RR, NG

Writing and analysis: All authors

